# Acarbose improves cognitive function in a mouse model of normal aging but not Alzheimer’s disease

**DOI:** 10.64898/2026.04.28.721469

**Authors:** Shannon J. Moore, Geoffrey G. Murphy

## Abstract

**INTRODUCTION:** Declines in function occur in both “normal” aging (in the absence of disease) and age-related pathological contexts, like Alzheimer’s disease (AD). Whether “anti-aging” interventions (that extend lifespan) also promote cognitive function in aging and AD remains unexplored.

**METHODS:** We assessed the effect of acarbose (1000 ppm from 4 months of age) on spatial learning and memory using the Morris water maze in young adult (6 mo), mid-aged (12 mo), or aged (24 mo) cohorts of normal aging (Ntg-HET3) and AD-relevant (5xFAD-HET3) genetically heterogeneous mice.

**RESULTS:** In mid-aged and aged Ntg-HET3 mice, acarbose treatment resulted in performance equivalent to young adults. Conversely, acarbose failed to ameliorate age-related deficits in 5xFAD-HET3 mice.

**DISCUSSION:** This work demonstrates that anti-aging interventions can also promote cognitive longevity in normal aging. Further, it reinforces that AD is not simply accelerated aging and requires therapies beyond anti-aging interventions that target its unique molecular and cellular drivers.

## 1. Background/Introduction

Aging is associated with a decline in both physical health and cognitive function, significantly decreasing the quality of life for the affected individuals, their families, and caregivers. In recent years, the number of Americans over the age of 65 has increased considerably, and is projected to continue to rise rapidly in the coming decades [1].

Coupled with the overall increase in life expectancy [2], more and more people are living longer at older ages, magnifying the impact of age-related dysfunction on an increasingly large segment of society.

In addition, advancing age is the primary risk factor – far greater than smoking, high cholesterol, or lack of exercise – for developing nearly every chronic disease, including cancer, cardiovascular disease, arthritis, and Alzheimer’s disease (AD) [3]. This observation has led to the “geroscience hypothesis”, which posits that interventions that target the aging process could ameliorate a wide range of deficits and illnesses simultaneously and in parallel, producing broader benefits than disease-specific treatments alone [4].

To this end, in 2003 the National Institute on Aging launched the Interventions Testing Program (ITP) to evaluate drugs, nominated by members of the scientific community, for their “anti-aging” potential [5, 6]. This program has successfully identified a number of putative “anti-aging” interventions, using the primary outcome of lifespan extension in a genetically heterogeneous mouse line (UM-HET3) as a reflection of delayed aging.

Importantly, subsequent studies on these successful lifespan-extending drugs have demonstrated improvements in various physiological aspects of aging, including liver and cardiac function [7, 8] and reduced inflammatory response [9–12], suggesting that these candidates may indeed slow multiple deleterious aspects of the aging process. However, the effect of these interventions on cognitive function - a crucial aspect of aging - has remained largely understudied. Both “normal” non-pathological aging (in the absence of disease) and age-related neurodegenerative diseases like AD [13, 14] are significantly associated with substantial cognitive deficits with more than 40% of individuals over the age of 65 experiencing some form of memory impairment [15, 16]. Thus, it is essential to elucidate and integrate the effects of anti-aging interventions on cognitive longevity with those on lifespan and physiological function to identify and prioritize interventions that synergistically impart the most impactful benefit in the context of both aging and disease.

Among the compounds identified by the ITP, acarbose (ACA) is of particular interest because it has an excellent safety profile [17] and is already FDA-approved (marketed as Precose® by Bayer for the treatment of type 2 diabetes), well-positioning it to be rapidly repurposed as a potential anti-aging therapeutic. ACA acts by inhibiting a-glucosidases in the intestines, thereby slowing the breakdown of complex carbohydrates. It was initially nominated because it produces physiological effects similar to caloric restriction (CR), which is the most consistent and robust intervention associated with extending lifespan in both animals and humans [18–20]. Specifically, both ACA and CR reduce body weight and body fat and, importantly, improve age-related glucose dysregulation and blunt post-prandial spikes in blood glucose, which have previously been implicated in aging [21]. And, indeed, ACA treatment started at 4 months of age successfully extended lifespan in both male and female UM-HET3 mice [22].

Taken together, ACA represents an attractive candidate to evaluate the efficacy of an anti-aging drug on cognitive function across the lifespan in “normal” aging and in the context of age-related neurodegenerative disease. To this end, we generated an experimental mouse population by modifying the four-way cross breeding scheme established by the ITP to produce genetically heterogeneous UM-HET3 mice where half of the mice carry the 5xFAD transgenes (“5xFAD-HET3”), which contain five distinct genetically heritable mutations that have been identified in human families with early-onset Alzheimer’s disease (AD) [23], while the remaining half do not (“Ntg-HET3”) (see Methods section “2.1 Mice” for details). We then assessed cognitive function at three different timepoints in individual cohorts of mice using a cross-sectional design. Cohorts of Ntg-HET3 mice (reflecting a “normal aging” population) and 5xFAD-HET3 mice (reflecting an AD-relevant population) were maintained on standard chow (“control”) or given ACA-containing chow (1000 ppm) starting at 4 months of age. Spatial learning and memory were evaluated using the Morris water maze (MWM) in young adult (6 mo), middle-aged (12 mo), and aged (24 mo) mice. We found that ACA effectively promoted cognitive function in normal aging, with an age-dependent improvement in spatial memory performance in Ntg-HET3 from 6 to 24 months but failed to ameliorate cognitive impairments that emerge with age in the 5xFAD-HET3 mice.

## 2. Methods

### 2.1 Mice

All procedures were approved by and performed in accordance with the Institutional Animal Care and Use Committee (IACUC) at the University of Michigan. Experimental mice were generated in house by following a modified four-way crossbreeding scheme based on that described in the NIA-funded ITP [5, 6] (see **Figure 1A**). In the ITP design, four well-characterized inbred isogenic strains, BALB/cByJ (CBy), C57BL/6J (B6), C3H/HeJ (C3), DBA/2J (D2), are used as founders to produce two distinct F1 hybrid lines: female CBy crossed with male B6 generate CByB6F1 mice and female C3 crossed with male D2 produce C3D2F1 mice. Then, a subsequent cross between female CByB6F1 and male C3D2F1 produces experimental UM-HET3 mice. As a result, each UM-HET3 mouse is genetically unique, containing a distinct, randomized combination of 25% of the genes from each of the original founder lines.

**Figure 1.**
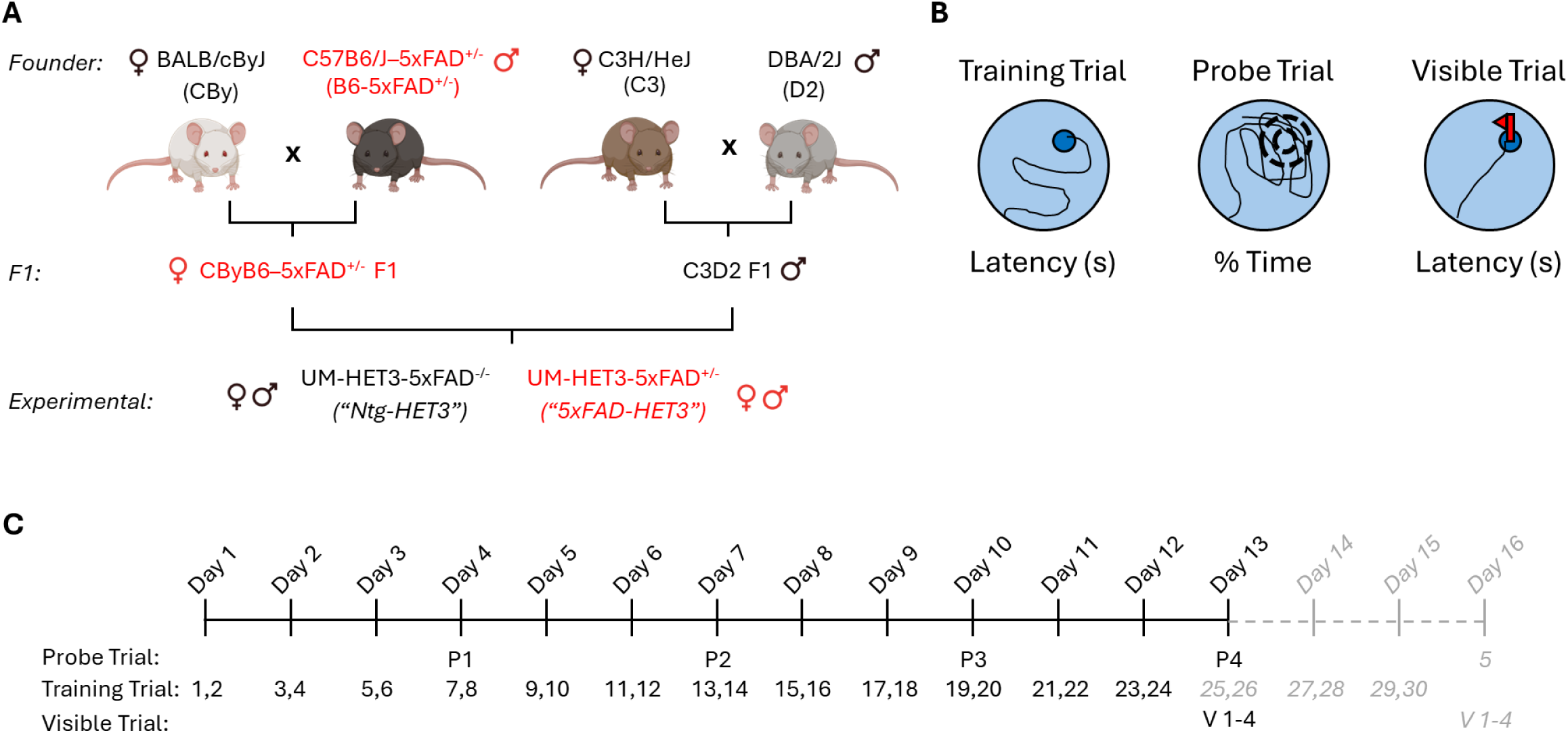
Experimental paradigm to investigate the effects of ACA on cognitive function in genetically heterogenous mouse models of “normal” aging (Ntg-HET3) and AD-relevant (5xFAD-HET3) phenotypes. (*A*) To model heterogeneity in the human population, we used an established four-way cross breeding scheme, which incorporates four grandparental inbred mouse lines: BALB/cJ (Cby), C57BL/6J (B6), C3H/HeJ (C3), and DBA/2J (D2). We modified this design by using the well-known C57BL/6J-5xFAD (B6-5xFAD^+/-^) mouse in place of the standard B6 mouse to introduce five separate genetically heritable mutations that have been identified in human families with early-onset Alzheimer’s disease (AD). Together, this results in an experimental population where half of the individuals are “wild-type” UM-HET3 mice (Ntg-HET3) and the other half harbor AD-associated mutations (5xFAD-HET3). (*B*) Cognitive function was assessed using the Morris Water Maze (MWM) in which mice are trained to use distal spatial cues to learn and remember the location of an escape platform. White lines indicate a representative swim paths. (*B, left*) During training trials (when the platform is present and submerged so that it is not visible, as indicated by the small dark blue circle), learning is reflected by lower latencies to reach the platform. (*B, middle*) Memory for the platform location is tested in probe trials (with the platform removed) and reflected by a spatially selective search strategy targeting the area where the platform was previously located (in these experiments, the target area comprised a circle with a 15 cm radius, centered on the platform’s previous location as shown by the dashed concentric circles). (*B, right*) Finally, visible trials (where the submerged platform is marked by a distinct, highly visible proximal cue, as shown by the small dark blue circle and red flag) are conducted to assess any sensory, motor, or motivational deficits that could confound the interpretation of learning and memory performance. (*C*) Experimental timeline for MWM. To learn the platform location, all mice were given 2 training trials per day for 12 days (except for the aged 5xFAD-HET3 mice, who required additional training; grey labels and dashed line); 24 hours after the last training trial in every 3-day period, a probe trial was conducted to assess memory performance (P1 – P4; aged 5xFAD-HET3 mice received an additional probe trial (P5) after their final set of 3 training days). After the final probe trial (P4 for all mice except P5 for aged 5xFAD-HET3), 4 visible platform trials were conducted.

We modified the ITP breeding scheme by replacing the isogenic inbred B6 founder with 5xFAD mice, which are also maintained on a B6 background. The B6-5xFAD line incorporates two hemizygous transgenes at a single insertion site whose expression is regulated by neural-specific elements of the mouse Thy1 promoter. These transgenes encode the human amyloid precursor protein (APP) and presenilin1 (PS1), containing five total causal mutations that were identified in familial cohorts with early onset AD (three in APP: Swedish (K670N, M671L), Florida (I716V), and London (V717I); and two in PS1: M146L and L286V). The B6-5xFAD line is a well-characterized mouse model of AD-relevant phenotypes, including beta amyloid plaque neuropathology and age-dependent deficits in cognitive function [23]. Thus, crossing female CBy mice (RRID:IMSR_JAX:000651) with male B6-5xFAD mice (RRID:MMRRC_034848-JAX) produces CByB6 F1 offspring where half inherit the 5xFAD transgenes (CByB6-5xFAD^+/-^). Then, by crossing these 5xFAD-inheriting F1 females with male C3D2 F1 mice (RRID:IMSR_JAX:100004), we produced our experimental population, which consisted of sibling cage mates (male and female) where with half of the mice carry the 5xFAD transgene (“5xFAD-HET3”) and half do not (“Ntg-HET3”; equivalent to “wild-type” UM-HET3 used in ITP studies). All mice needed for breeding were purchased from The Jackson Laboratory (Bar Harbor, MA).

### 2.2 Experimental Design: ACA Administration and Experimental Cohorts

In alignment with studies conducted by the ITP evaluating the effect of ACA on lifespan [22], at 4 mo of age, mice were pseudo-randomly divided into two groups, one of which was maintained on standard chow (“control”) and one of which began receiving ACA-compounded chow (1000 ppm), available *ad libitum* (all chow from TestDiet, Richmond, IN). Chow was assigned on a per-cage basis, which was maintained throughout the entire experimental timeline. Cross-sectional cohorts of mice were assessed for cognitive function using the MWM at three different ages: young adult (6 mo), middle-aged (12 mo) and aged (24 mo). All cohorts were initially balanced for both sex and genotype (presence or absence of the 5xFAD transgene), but due to unexpectedly high rates of aggression in male cages, significant loss due to attrition reduced the power to explicitly examine the effect of sex. For all analyses presented herein, males and females were collapsed within each age x treatment group.

### 2.3 Morris water maze (MWM)

The MWM was used to evaluate learning and memory, based on protocols previously described [24, 25] (see **Figure 1**). The pool was 120 cm in diameter with a removable platform that was 10 cm in diameter and placed in a room with distal extra-maze cues (e.g. high contrast posters placed on the walls); the water was maintained at ∼22-25’C and made opaque using non-toxic white paint. All trials were video recorded with a camera mounted directly above the pool and connected to a computer running WaterMaze software (Actimetrics, Wilmette, IL) for data acquisition, storage, and analysis.

#### 2.3.1 Spatial Learning/Training Trials

Mice were trained to learn the location of a submerged (hidden) platform (located ∼1 cm below the surface of the water) in two trials/day (back-to-back for an individual mouse with a 10-second intertrial interval). Prior to each training trial, mice were put on the platform for ∼10 seconds and then placed in the pool facing the pool wall at pseudo-random start locations (same for all mice on each day). Mice were allowed to swim until they found the platform, up to a maximum of 60 seconds, which was recorded as “latency to platform”. A reduction in latency across days of training was taken to reflect successful spatial learning. Mice that did not find the platform within 60 seconds were gently guided to the platform by the experimenter.

#### 2.3.2 Spatial Memory Tests/Probe Trials

Mice were tested for memory of the platform location in probe trials (during which the submerged platform was removed from the pool) interspersed throughout training trials; one probe trial was given after each set of three training days (6 training trials), ∼24 hours after the last training trial (on the previous day). For each probe trial, mice were placed in the pool facing the wall directly opposite from where the platform was located in the training trials and allowed to swim for 60 seconds before being removed from the pool. During each probe trial, time spent searching target area (defined as a 30-cm diameter circle, centered on where the platform was located) was measured. This stringent definition for the target area was chosen to ensure robust detection of a highly selective spatial search strategy. Improved performance over the course of the experiment was reflected by an increase in the amount of time spent searching the target area across probe trials. In addition, to ensure that the search strategy was selective for the target location, we compared the percent time spent in the target area on each probe trial to that which would be expected by “chance” (with a random or non-spatially selective search strategy), which was calculated as the ratio of the circular target area divided by the total pool area (225 cm^2^/3600 cm^2^ = 6.25%). Performance significantly greater than “chance” was taken to indicate successful formation of a spatial memory for the platform location.

#### 2.3.3 Visible Platform Trials

At the end of the experiment, immediately after the final probe trial, visible platform trials where the submerged platform was marked by a distinct proximal cue (flag on the platform) were conducted to determine whether there were any differences in non-cognitive functions (e.g. visual or motor impairments) that could confound the interpretation of learning and memory performance. Four total visible platform trials were conducted in two sets of two back-to-back trials, with the marked submerged platform moved to a new location between each set of two trials. Prior to each visible trial, mice were put on the platform for ∼10 seconds and then placed in the pool facing the pool wall at pseudo-random start locations (same for all mice on each day). Mice were allowed to swim until they found the platform, up to a maximum of 60 seconds, which was recorded as “latency to platform”.

### 2.4 Analysis

Metrics to assess spatial learning (latency to reach platform), spatial memory (percent time in target area), and motor performance (swim speed) were calculated and exported automatically via WaterMaze software (Actimetrics, Wilmette, IL). All data were analyzed using Prism (GraphPad, San Diego, CA) with 3-factor or 2-factor repeated measures ANOVA and Bonferroni’s post-hoc tests or Welsh’s corrected t-tests; full statistical details are reported in Tables 1-4. Any animal failing to meet specified criteria [26] was excluded from all analysis (24 out of 277 mice); these exclusion criteria were: 1. Died during the experiment; 2. Failed to reach the visible platform on all four trials (i.e. visible platform trial latency = 60 s); 3. Floating during two or more probe trials (reflected by a swim speed less than half of that age x genotype group’s average); 4.Severe thigmotaxis on two or more probe trials (reflected by percent time near walls >40%).

**Table 1.** Descriptive Statistics by Group. Ntg-HET3. Table 1. Descriptive Statistics by Group. Table 1 shows mean +/- SEM for the following parameters for all treatment x age cohorts: n (M,F) is the number of male and female mice in each group; Weight (g) was measured for all mice at the start of the Morris water maze; Swim speed (cm/s) was averaged across all relevant trials (training or probe, as indicated) for an individual mouse, then averaged by group; Latency to Visible Platform (s) was averaged across 4 visible platform trials for an individual mouse, then averaged by group. A within-age group comparison between control- and ACA-treated was performed for each parameter using Welch’s t-test; statistically significant differences are bolded and indicated by an asterisk.

|  | n (M,F) | Weight (g) | Swim Speed - Training (cm/s) | Swim Speed - Probe (cm/s) | Latency to Visible Platform (s) |
| --- | --- | --- | --- | --- | --- |
| Control | 20 (7M, 13F) | 36.5 ± 1.6 | 16.5 ± 0.5 | 21.5 ± 0.6 | 6.1 ± 0.3 |
| ACA | 25 (13M, 12F) | 30.6 ± 1.0 | 16.8 ± 0.5 | 22.2 ± 0.6 | 6.5 ± 0.8 |
| Welch's t-test |  |  |  |  |  |
| p value (t, df) |  | *p < 0.01<br>(3.2, 32.78) | p = 0.63<br>(0.48, 43.0) | p = 0.46<br>(0.75, 42.2) | p = 0.59<br>(0.54, 31.2) |

**Middle-aged (12 mo)**
|  | n (M,F) | Weight (g) | Swim Speed - Training (cm/s) | Swim Speed - Probe (cm/s) | Latency to Visible Platform (s) |
| --- | --- | --- | --- | --- | --- |
| Control | 20 (7M, 13F) | 43.7 ± 1.5 | 15.1 ± 0.7 | 18.5 ± 0.6 | 8.7 ± 0.8 |
| ACA | 26 (10M, 16F) | 35.8 ± 1.0 | 16.0 ± 0.4 | 19.8 ± 0.6 | 7.8 ± 0.7 |
| Welch's t-test |  |  |  |  |  |
| p value (t, df) |  | *p<0.001<br>(3.9, 25.8) | p=0.30<br>(1.1, 30.1) | p=0.14<br>(1.5, 42.8) | p=0.42<br>(0.8, 40.6) |

**Aged (24 mo)**
|  | n (M,F) | Weight (g) | Swim Speed - Training (cm/s) | Swim Speed - Probe (cm/s) | Latency to Visible Platform (s) |
| --- | --- | --- | --- | --- | --- |
| Control | 23 (10M, 13F) | 39.3 ± 1.2 | 13.3 ± 0.5 | 16.6 ± 0.6 | 19.6 ± 2.3 |
| ACA | 25 (8M, 17F) | 34.2 ± 1.1 | 13.6 ± 0.5 | 17.6 ± 0.6 | 17.8 ± 2.6 |
| Welch's t-test |  |  |  |  |  |
| p value (t, df) |  | *p<0.01<br>(3.0, 45.0) | p=0.63<br>(0.48, 45.6) | p=0.24<br>(1.2, 46.0) | p=0.61<br>(0.51, 44.5) |

**5xFAD-HET3****Young adult (6 mo)**
|  | n (M,F) | Weight (g) | Swim Speed - Training (cm/s) | Swim Speed - Probe (cm/s) | Latency to Visible Platform (s) |
| --- | --- | --- | --- | --- | --- |
| Control | 21 (9M, 12F) | 37.4 ± 1.6 | 17.5 ± 0.6 | 23.1 ± 0.7 | 7.1 ± 0.5 |
| ACA | 22 (13M, 9F) | 30.9 ± 0.8 | 17.7 ± 0.4 | 23.7 ± 0.6 | 5.9 ± 0.5 |
| Welch's t-test |  |  |  |  |  |
| p value (t, df) |  | *p<0.01<br>(3.6, 29.8) | p=0.75<br>(0.33, 37.3) | p=0.50<br>(0.69, 40.1) | p=0.12<br>(1.6, 40.6) |

**Middle-aged (12 mo)**
|  | n (M,F) | Weight (g) | Swim Speed - Training (cm/s) | Swim Speed - Probe (cm/s) | Latency to Visible Platform (s) |
| --- | --- | --- | --- | --- | --- |
| Control | 15 (6M, 9F) | 40.8 ± 1.5 | 16.6 ± 0.5 | 20.2 ± 0.7 | 9.5 ± 1.0 |
| ACA | 22 (11M, 11F) | 33.5 ± 1.0 | 17.0 ± 0.5 | 21.4 ± 0.6 | 9.8 ± 0.6 |
| Welch's t-test |  |  |  |  |  |
| p value (t, df) |  | *p<0.0001<br>(4.5, 34.0) | p=0.51<br>(0.66, 34.0) | p=0.20<br>(1.3, 29.8) | p=0.77<br>(0.3, 25.2) |

**Table 1. Descriptive Statistics by Group.**
|  | n (M,F) | Weight (g) | Swim Speed - Training (cm/s) | Swim Speed - Probe (cm/s) | Latency to Visible Platform (s) |
| --- | --- | --- | --- | --- | --- |
| Control | 17 (7M, 10F) | 26.3 ± 1.3 | 15.9 ± 0.5 | 18.5 ± 0.6 | 31.2 ± 4.0 |
| ACA | 18 (4M, 14F) | 26.6 ± 1.6 | 15.3 ± 0.4 | 18.0 ± 0.7 | 27.0 ± 3.2 |
| Welch's t-test |  |  |  |  |  |
| p value (t, df) |  | p=0.88<br>(0.15, 32.4) | p=0.36<br>(0.94, 31.8) | p=0.58<br>(0.55, 32.4) | p=0.42<br>(0.83, 31.2) |

**Table 2.**
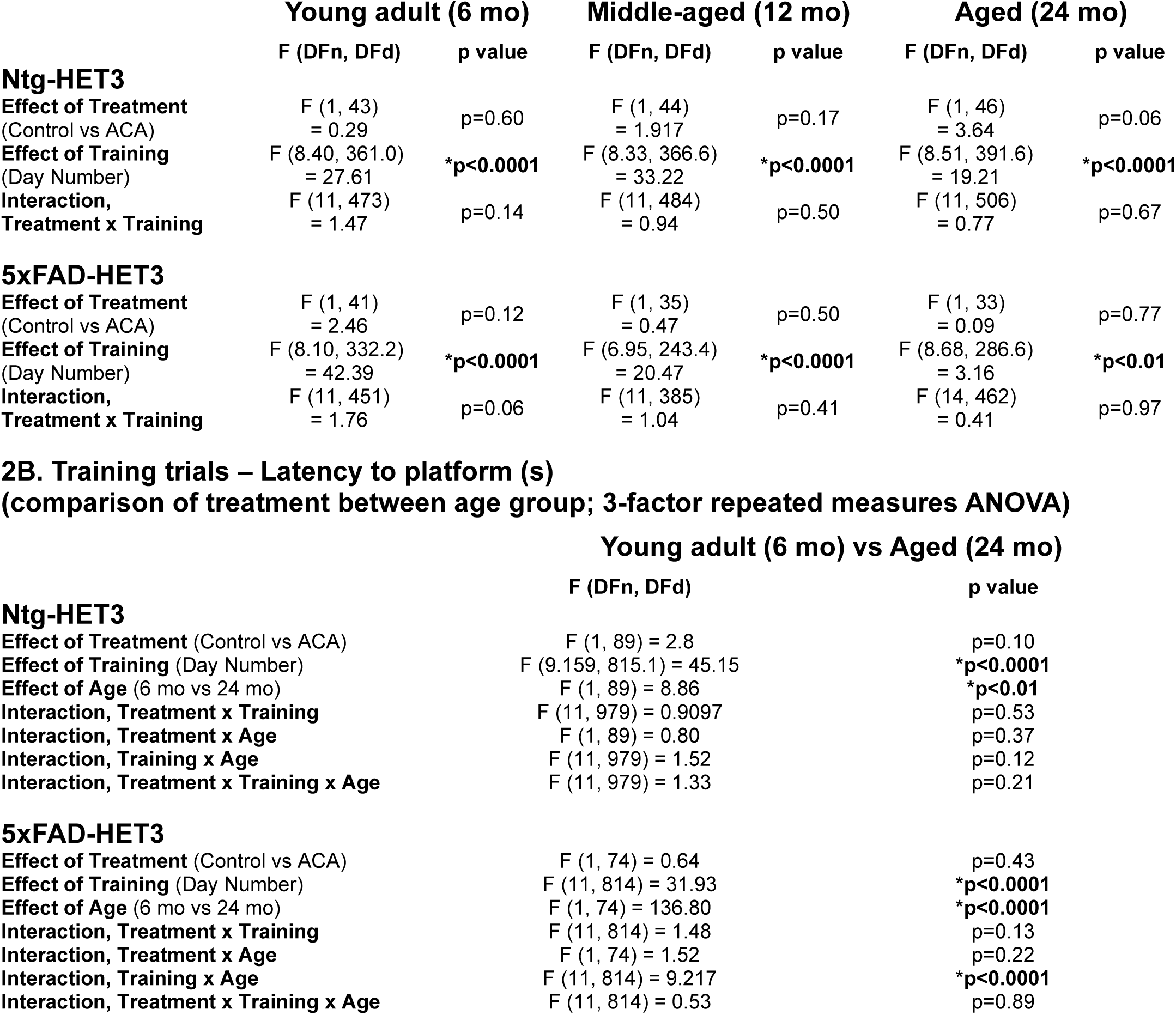
Training Trial Statistics by Group. 2A. Training trials – Latency to platform (s) Table 2. Training Trial Statistics by Group. Table 2A shows the results from a 2-factor (treatment and day of training) repeated measures ANOVA of latency to platform (s) between control- and ACA-treated groups within each age cohort. Table 2B shows the results from a 3-factor (treatment, day of training, and age) repeated measures ANOVA of latency to platform (s) between control- and ACA-treated groups between the young adult and aged cohort. For both tests, statistically significant differences are bolded and indicated by an asterisk.

**(comparison of treatment within age group; 2-factor repeated measures ANOVA)**
|  | **Young adult (6 mo)** |  | **Middle-aged (12 mo)** |  | **Aged (24 mo)** |  |
| --- | --- | --- | --- | --- | --- | --- |
|  | **F (DFn, DFd)** | **p value** | **F (DFn, DFd)** | **p value** | **F (DFn, DFd)** | **p value** |
| **Ntg-HET3** |  |  |  |  |  |  |
| **Effect of Treatment**<br>(Control vs ACA) | F (1, 43)<br>= 0.29 | p=0.60 | F (1, 44)<br>= 1.917 | p=0.17 | F (1, 46)<br>= 3.64 | p=0.06 |
| **Effect of Training**<br>(Day Number) | F (8.40, 361.0)<br>= 27.61 | **\*p<0.0001** | F (8.33, 366.6)<br>= 33.22 | **\*p<0.0001** | F (8.51, 391.6)<br>= 19.21 | **\*p<0.0001** |
| **Interaction,**<br>**Treatment x Training** | F (11, 473)<br>= 1.47 | p=0.14 | F (11, 484)<br>= 0.94 | p=0.50 | F (11, 506)<br>= 0.77 | p=0.67 |
| **5xFAD-HET3** |  |  |  |  |  |  |
| **Effect of Treatment**<br>(Control vs ACA) | F (1, 41)<br>= 2.46 | p=0.12 | F (1, 35)<br>= 0.47 | p=0.50 | F (1, 33)<br>= 0.09 | p=0.77 |
| **Effect of Training**<br>(Day Number) | F (8.10, 332.2)<br>= 42.39 | **\*p<0.0001** | F (6.95, 243.4)<br>= 20.47 | **\*p<0.0001** | F (8.68, 286.6)<br>= 3.16 | **\*p<0.01** |
| **Interaction,**<br>**Treatment x Training** | F (11, 451)<br>= 1.76 | p=0.06 | F (11, 385)<br>= 1.04 | p=0.41 | F (14, 462)<br>= 0.41 | p=0.97 |

**Table 2. Training Trial Statistics by Group.**
|  | **F (DFn, DFd)** | **p value** |
| --- | --- | --- |
| **Ntg-HET3** |  |  |
| **Effect of Treatment** (Control vs ACA) | F (1, 89) = 2.8 | p=0.10 |
| **Effect of Training** (Day Number) | F (9.159, 815.1) = 45.15 | **\*p<0.0001** |
| **Effect of Age** (6 mo vs 24 mo) | F (1, 89) = 8.86 | **\*p<0.01** |
| **Interaction, Treatment x Training** | F (11, 979) = 0.9097 | p=0.53 |
| **Interaction, Treatment x Age** | F (1, 89) = 0.80 | p=0.37 |
| **Interaction, Training x Age** | F (11, 979) = 1.52 | p=0.12 |
| **Interaction, Treatment x Training x Age** | F (11, 979) = 1.33 | p=0.21 |
| **5xFAD-HET3** |  |  |
| **Effect of Treatment** (Control vs ACA) | F (1, 74) = 0.64 | p=0.43 |
| **Effect of Training** (Day Number) | F (11, 814) = 31.93 | **\*p<0.0001** |
| **Effect of Age** (6 mo vs 24 mo) | F (1, 74) = 136.80 | **\*p<0.0001** |
| **Interaction, Treatment x Training** | F (11, 814) = 1.48 | p=0.13 |
| **Interaction, Treatment x Age** | F (1, 74) = 1.52 | p=0.22 |
| **Interaction, Training x Age** | F (11, 814) = 9.217 | **\*p<0.0001** |
| **Interaction, Treatment x Training x Age** | F (11, 814) = 0.53 | p=0.89 |

**Table 3.** Probe Trial Statistics by Group. 3A. Probe trials - Time in Target Area (%) Table 3. Probe Trial Statistics by Group. Table 3A shows the results from a 2-factor (treatment and day of training) repeated measures ANOVA of percent time in target area between control- and ACA-treated groups within each age cohort. Table 3B shows the results from a 3-factor (treatment, day of training, and age) repeated measures ANOVA of percent time in target area between control- and ACA-treated groups between the young adult and aged cohort. For both tests, statistically significant differences are bolded and indicated by an asterisk.

|  | Young adult (6 mo) |  | Middle-aged (12 mo) |  | Aged (24 mo) |  |
| --- | --- | --- | --- | --- | --- | --- |
|  | F (DFn, DFd) | p value | F (DFn, DFd) | p value | F (DFn, DFd) | p value |
| **Ntg-HET3** |  |  |  |  |  |  |
| Effect of Treatment (Control vs ACA) | F (1, 43) = 0.11 | p=0.74 | F (1, 44) = 1.24 | p=0.27 | F (1, 46) = 5.18 | **\*p=0.03** |
| Effect of Training (Day Number) | F (2.93, 126.0) = 33.63 | **\*p<0.0001** | F (2.40, 105.6) = 35.94 | **\*p<0.0001** | F (2.28, 105.0) = 29.11 | **\*p<0.0001** |
| Interaction, Treatment x Training | F (3, 129) = 1.40 | p=0.25 | F (3, 132) = 2.94 | **\*p=0.04** | F (3, 138) = 1.83 | p=0.14 |
| **5xFAD-HET3** |  |  |  |  |  |  |
| Effect of Treatment (Control vs ACA) | F (1, 41) = 0.44 | p=0.51 | F (1, 35) = 2.17 | p=0.15 | F (1, 33) = 0.53 | p=0.47 |
| Effect of Training (Day Number) | F (2.89, 118.3) = 22.40 | **\*p<0.0001** | F (2.41, 84.16) = 23.31 | **\*p<0.0001** | F (3.11, 102.5) = 11.39 | **\*p<0.0001** |
| Interaction, Treatment x Training | F (3, 123) = 0.33 | p=0.80 | F (3, 105) = 0.13 | p=0.94 | F (4, 132) = 1.44 | p=0.22 |

**Table 3. Probe Trial Statistics by Group.**
|  | F (DFn, DFd) | p value |
| --- | --- | --- |
| **Ntg-HET3** |  |  |
| Effect of Treatment (Control vs ACA) | F (1, 120) = 4.76 | **\*p=0.03** |
| Effect of Training (Day Number) | F (3, 360) = 76.51 | **\*p<0.0001** |
| Effect of Age (6 mo vs 24 mo) | F (1, 120) = 0.38 | p=0.54 |
| Interaction, Treatment x Training | F (3, 360) = 0.78 | p=0.51 |
| Interaction, Treatment x Age | F (1, 120) = 2.23 | p=0.14 |
| Interaction, Training x Age | F (3, 360) = 0.06 | p=0.98 |
| Interaction, Treatment x Training x Age | F (3, 360) = 2.84 | **\*p=0.04** |
| **5xFAD-HET3** |  |  |
| Effect of Treatment (Control vs ACA) | F (1, 74) = 0.02 | p=0.89 |
| Effect of Training (Day Number) | F (3, 222) = 31.20 | **\*p<0.0001** |
| Effect of Age (6 mo vs 24 mo) | F (1, 74) = 27.92 | **\*p<0.0001** |
| Interaction, Treatment x Training | F (3, 222) = 1.751 | p=0.16 |
| Interaction, Treatment x Age | F (1, 74) = 0.8125 | p=0.37 |
| Interaction, Training x Age | F (3, 222) = 2.086 | p=0.10 |
| Interaction, Treatment x Training x Age | F (3, 222) = 0.70 | p=0.55 |

**Table 4.**
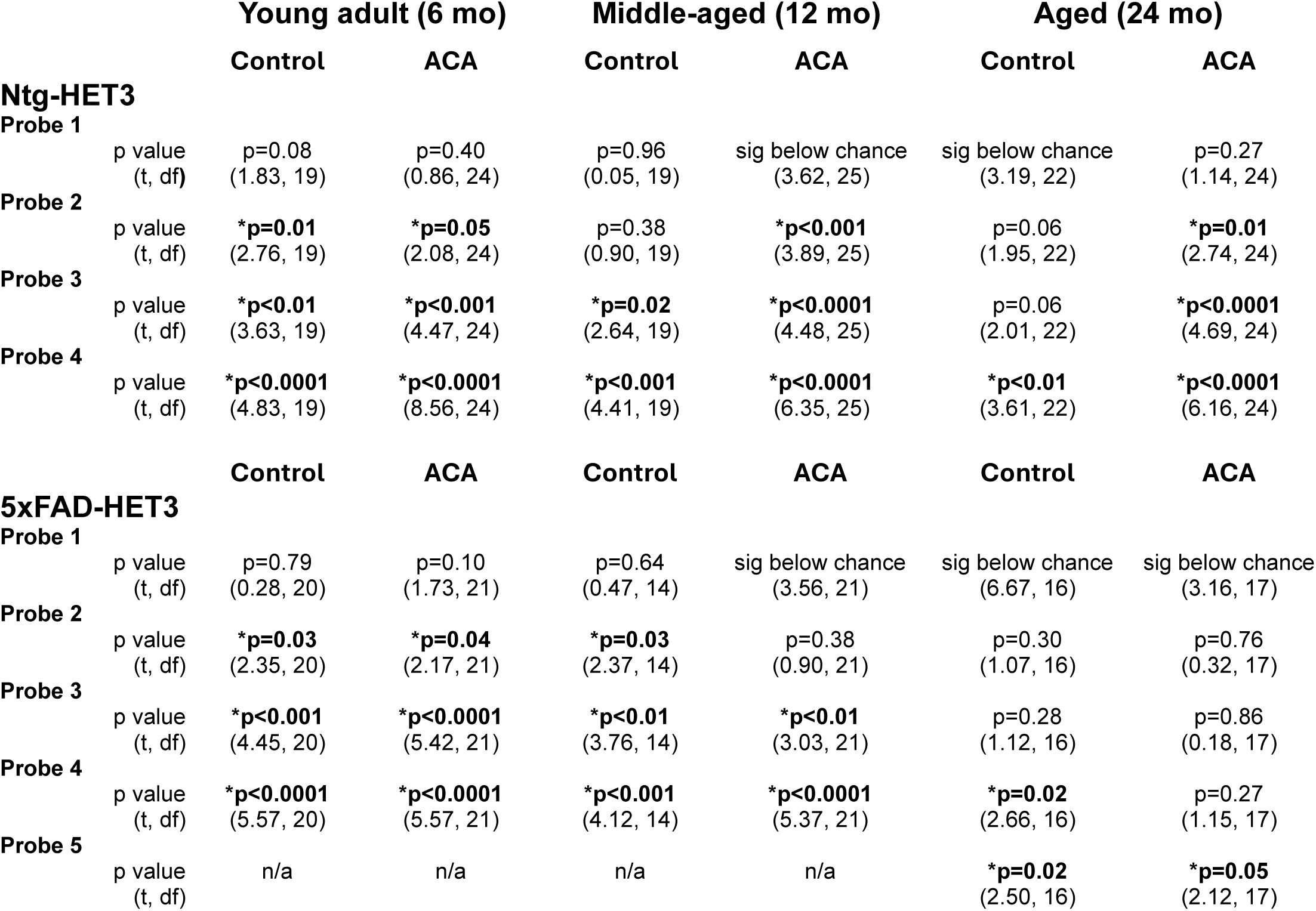
Probe Trial Statistics against “chance”. Probe Trial Statistics: Time in Target Area (%) (comparison to “chance”; Welch’s one-sample t-test) Table 4. Probe Trial Statistics against “chance”. Table 4 shows the results for each treatment x age group of a Welch’s one-sample t-test for each probe trial against “chance” performance (calculated as the ratio of the target area to the pool area, see Methods 2.3.2 Spatial Memory Tests/Probe Trials for details). Statistically significant differences are bolded and indicated by an asterisk.

## 3. Results

### 3.1 The effect of ACA in Ntg-HET3 mice, reflecting a “normal aging” population

To determine the effect of ACA treatment on cognitive performance across the lifespan in a normal aging population (in the absence of disease-related changes), male and female Ntg-HET3 mice were either given with ACA-containing chow (1000 ppm) starting at 4 months of age or maintained on standard chow (“control”). All mice were weighed prior to beginning cognitive assessment; consistent with previous reports [8, 22], ACA-treated mice at all ages weighed significantly less than their control-treated counterparts **(Table 1)**. Importantly, the observed reduction in body weight demonstrates that ACA was successfully administered and metabolized, exerting its expected pharmacological effects. Therefore, any absence of a cognitive effect with ACA treatment is unlikely to be due to a failure of drug delivery or efficacy.

Control and ACA-treated mice were behaviorally characterized in cohorts of young adult (6 month), middle-aged (12 month), or aged (24 month) mice in a cross-sectional design. Cognitive performance was assessed using the MWM, a hippocampus-dependent task in which mice use visual cues located around the pool to learn and remember the position of a submerged (hidden) platform. Importantly, swim speeds and latency to reach a visible platform (marked with a proximal cue) were also measured in these groups; there were no differences on either of these measures between control-and ACA-treated mice at any age, suggesting that the effect of ACA on learning and memory performance was not confounded by differences in sensory-motor abilities, motivation, or procedural capabilities **(Table 1)**.

### 3.2 Spatial learning is not improved by ACA in Ntg-HET3 mice

In all cohorts, control- and ACA-treated mice performed equally well on training trials; each group exhibited a significant reduction in latency to reach the platform across days of training (p<0.0001, effect of training in each cohort) and there were no significant differences in performance between control and ACA-treated mice in any age group (effect of treatment in: young adult, p=0.29; middle-aged, p=0.17; aged, p=0.06; **Figure 2A_1_-C_1_ and Table 2A**). These data suggest that both control- and ACA- treated mice at all ages showed some ability to learn the platform location, and that ACA did not significantly affect the rate or degree of learning within any cohort. However, comparing performance between cohorts revealed an age-related deficit (effect of age p<0.01; **Table 2B**), with aged mice exhibiting a significantly longer latency to reach the platform. Importantly, consistent with the lack of an ACA-mediated improvement in spatial learning at any age, the age-related deficit was not ameliorated by ACA treatment, reflected by the lack of an effect of treatment or interaction of treatment with age between cohorts (effect of: treatment, p=0.10; treatment x age, p=0.37; treatment x training x age, p=0.21; **Table 2B**).

**Figure 2.**
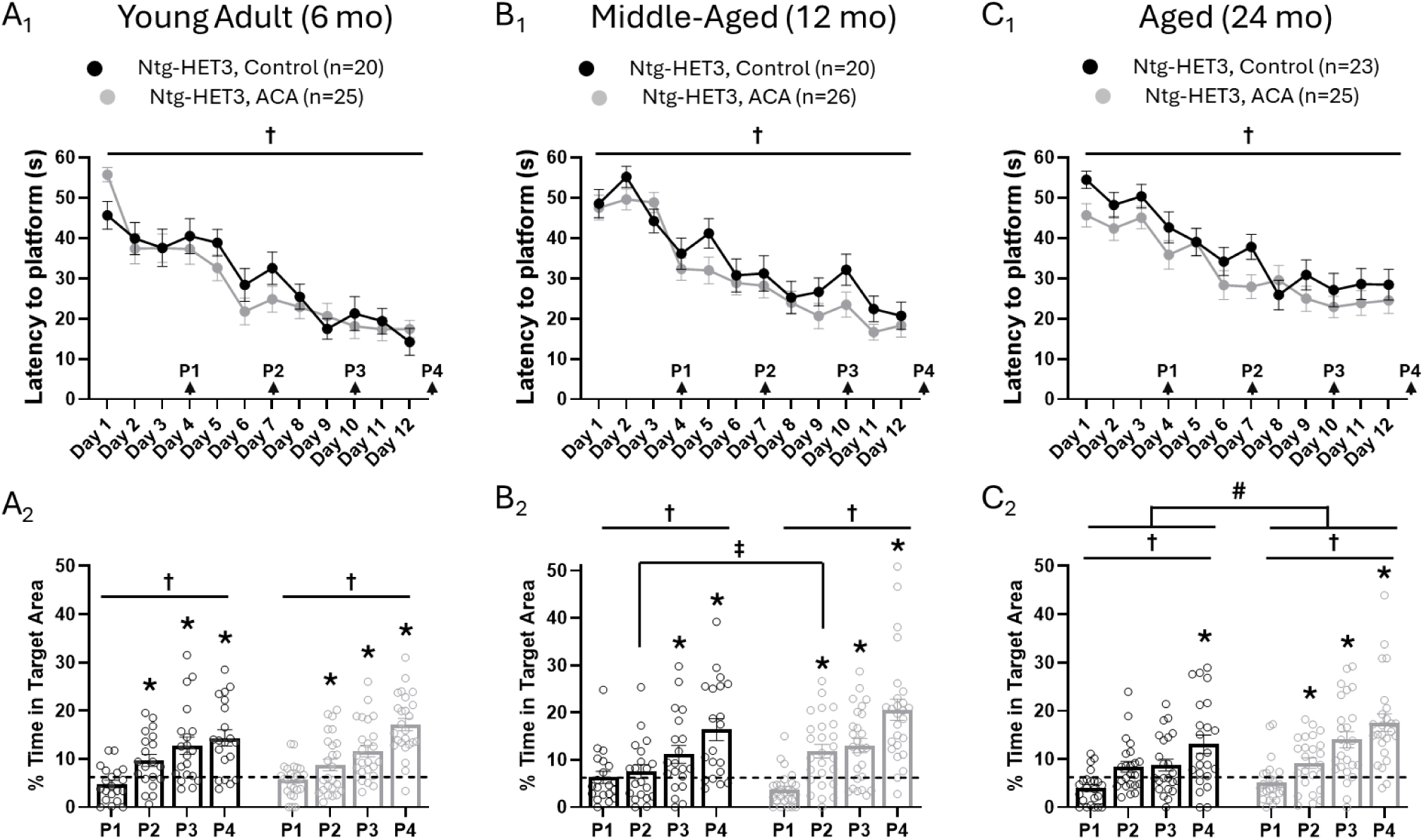
ACA age-dependently improves spatial memory in Ntg-HET3 mice. Separate groups of Ntg-HET3 mice were treated with ACA chow (1000 ppm, starting at 4 mo of age; shown in grey) or remained on control diet (“control”, shown in black) and assessed for hippocampus-dependent learning and memory using the MWM at young adult (6 mo), middle-aged (12 mo), or aged (24 mo) timepoints. **(*Top panels, A_1_-C_1_)*** Mice were given 2 trials/day over 12 days of training. Both control- and ACA mice in all age groups exhibited a significant reduction in latency to reach the platform over days of training († denotes effect of time, p<0.0001), with no difference in performance between groups at any age (effect of treatment in *A_1_*: young adult, p=0.60; *B_1_*: middle-aged, p=0.17; *C_1_*: aged, p=0.06). See Table 2 full statistical details. ***(Bottom panels, A_2_-C_2_)*** Four probe trials were interspersed throughout training at the times indicated by the arrows in A_1_-C_1_ (P1 = Probe 1, etc). *(A_2_-C_2_)* Both control- and ACA mice in all age groups demonstrated a spatially selective search strategy, as evidenced by a significant increase in the percent time spent searching the target area (circle with 15 cm radius, centered on the training platform’s location) over successive probe trials († denotes effect of time, p<0.0001). *(A_2_)* Both groups of mice exhibited equally good performance as young adults (effect of treatment in *A_2_*: young adult, p=0.74). *(B_2_-C_2_)* However, middle-aged and aged ACA-treated mice showed significantly better performance compared to control-treated mice in their age group (*B_2_*: middle-aged, ‡ denotes treatment x time interaction with Bonferroni post-hoc, p<0.05; *C_2_*: aged, **^#^** denotes effect of treatment, p<0.05). *(A_2_-C_2_)* Further, performance on each probe trial was compared to “chance” (indicated by the dashed line), which is the percent time an animal would be expected to be in the target area “by chance” if they exhibited a random or non-spatially selective search strategy; performance significantly above chance reflects the successful formation of a selective search strategy (* denotes p<0.05). *(A_2_)* In the young adult group, both control- and ACA-treated mice exhibited performance significantly above chance by Probe 2. *(B_2_)* However, in the middle-aged group, while ACA-treated mice still showed a spatially selective search strategy by Probe 2, this was not demonstrated in control-treated mice until Probe 3. *(C_2_)* Finally, in the aged group, ACA-treated animals again showed a spatially selective search strategy by Probe 2, but control-treated mice did not show performance significantly above chance until Probe 4. See Tables 3 and 4 for full statistical details.

### 3.3 Age-related deficits in spatial memory are ameliorated by ACA in Ntg-HET3 mice

In the young adult cohort, both control and ACA-treated groups exhibited equally good performance on probe trials, showing a significant increase in percent time spent in the target area over the course of the experiment (effect of training, p<0.0001; effect of treatment, p=0.74; **Figure 2A_2_**; **Table 3A**). In addition, when compared to “chance”, both control and ACA-treated mice spent significantly more time in the target area by Probe 2 (control-treated, p=0.01; ACA-treated, p=0.05), suggesting that mice in both groups had successfully formed a memory for the platform location by this point in the experiment (**Figure 2A_2_**; **Table 4**).

In the middle-aged cohort, both control and ACA-treated groups again showed significant increases in percent time spent in the target area over the course of the experiment and but there was also an interaction between treatment and probe trial number, suggesting that the rate of the improvement in performance was faster in ACA-treated mice (effect of training, p<0.0001; effect of treatment x training, p=0.04; **Figure 2B_2_**; **Table 3A**). Further, when examining performance on individual probe trials, ACA-treated mice exhibited a percent time in the target area significantly greater than “chance” by Probe 2 (control-treated, p=0.38; ACA-treated, p<0.001), while control-treated mice did not show performance above “chance” until Probe 3 (control-treated, p=0.02; ACA-treated, p<0.0001; **Figure 2B_2_**; **Table 4**). Taken together, these results suggest that ACA modestly but significantly improved spatial memory in middle-aged mice.

ACA had the most substantial effect on performance in the aged cohort. While both control and ACA-treated groups exhibited a significant increase in time spent in the target area over the course of the experiment, ACA-treated mice performed significantly better than control-treated mice across all probe trials (effect of training, p<0.0001; effect of treatment, p=0.03; **Figure 2C_2_**; **Table 3A**), suggesting they formed a better memory for the platform location. Further, when individual probe trials were examined, ACA-treated mice exhibited performance significantly above “chance” by Probe 2 (control-treated, p=0.06; ACA-treated, p=0.01), while an equivalent level of performance was not reached by control-treated mice until Probe 4 (control-treated, p<0.01; ACA-treated, p<0.0001; **Figure 2C**_2_; **Table 4**), suggesting that ACA robustly improved memory formation in aged mice.

Comparing performance between young and aged cohorts provided additional direct evidence that ACA successfully improved performance on probe trials in an age-dependent manner. Specifically, there was a significant three-way interaction between treatment, training (probe number), and age indicating that the increase in precent time spent in the target area over the course of the experiment in ACA-treated mice was age-specific (effect of treatment x training x age, p=0.04; **Table 3B**). This suggests that, while there was no effect of ACA treatment at a young age, its impact became increasingly pronounced with advancing age, from a modest enhancement in the rate of memory formation in middle-aged mice to a robust improvement in both the rate of formation and overall strength of spatial memory in aged mice. Taken together, these data suggest that ACA ameliorated age-related deficits in spatial memory, with the greatest impact at the oldest ages.

### 3.4 The effect of ACA in 5xFAD-HET3 mice, reflecting an AD-relevant population

Advancing age is the greatest risk factor for developing neurodegenerative diseases, including AD. Therefore, given the beneficial effects of ACA on cognitive function in a normal aging population, we next asked whether these benefits would extend to learning and memory impairments in a disease-relevant context. To address this question, we generated mice that inherited the 5xFAD transgenes, which harbor 5 distinct mutations identified in families with early onset AD [23], on a UM-HET3 background (“5xFAD-HET3”; see Methods). We treated both male and female 5xFAD-HET3 mice with ACA-containing chow (1000 ppm) starting at 4 months of age (while a “control” group was maintained on standard chow) and then behaviorally characterized mice using the MWM in young adult (6 month), middle-aged (12 month), or aged (24 month) cohorts in a cross-sectional design. Swim speeds and latency to reach a visible platform were also measured in these groups and showed no differences between control- and ACA-treated mice in each age group, suggesting that the effect of ACA on learning and memory performance was not confounded by differences in sensory-motor abilities, motivation, or procedural capabilities **(Table 1)**.

In both young adult and middle-aged cohorts, 5xFAD-HET3 mice treated with ACA weighed significantly less than their control-treated counterparts **(Table 1)**, consistent with expected effects of ACA administration. However, in the aged group, there was no significant difference in weight between control- and ACA-treated 5xFAD-HET3 mice, with both groups weighing notably less than their Ntg-HET3 counterparts (**Table 1**). This likely reflects disease-related weight loss in both the control- and ACA-treated mice, with a floor effect obscuring any further reduction due to ACA alone.

### 3.5 Spatial learning is not improved by ACA in 5xFAD-HET3 mice

In training trials, both control- and ACA-treated 5xFAD-HET3 mice within each cohort exhibited a significant reduction in the latency to reach the platform across days (effect of training in: young adult and middle-aged, p<0.0001 for each; aged, p<0.01) and showed no differences between treatment groups (effect of treatment in: young adult, p=0.12 ; middle-aged, p=0.50; aged, p=0.77), suggesting that all mice exhibited some degree of spatial learning for the platform location and that this performance was not affected by ACA treatment (**Figure 3A_1_-C_1_; Table 2A**). However, when performance was compared between cohorts, both control- and ACA-treated 5xFAD-HET3 mice in the aged group took significantly longer to reach the platform than young adult mice, indicating that there was an age-related deficit in spatial learning (effect of age, p<0.0001; **Table 2B**). Consistent with the lack of an effect of ACA within the aged cohort, there was no effect of treatment or interactions of treatment with age between cohorts (effect of: treatment, p=0.43; treatment x age, p=0.22; treatment x training x age, p=0.89; **Table 2B**), indicating that ACA does not improve spatial learning at any age or ameliorate age-related deficits in spatial learning in 5xFAD-HET3 mice.

**Figure 3.**
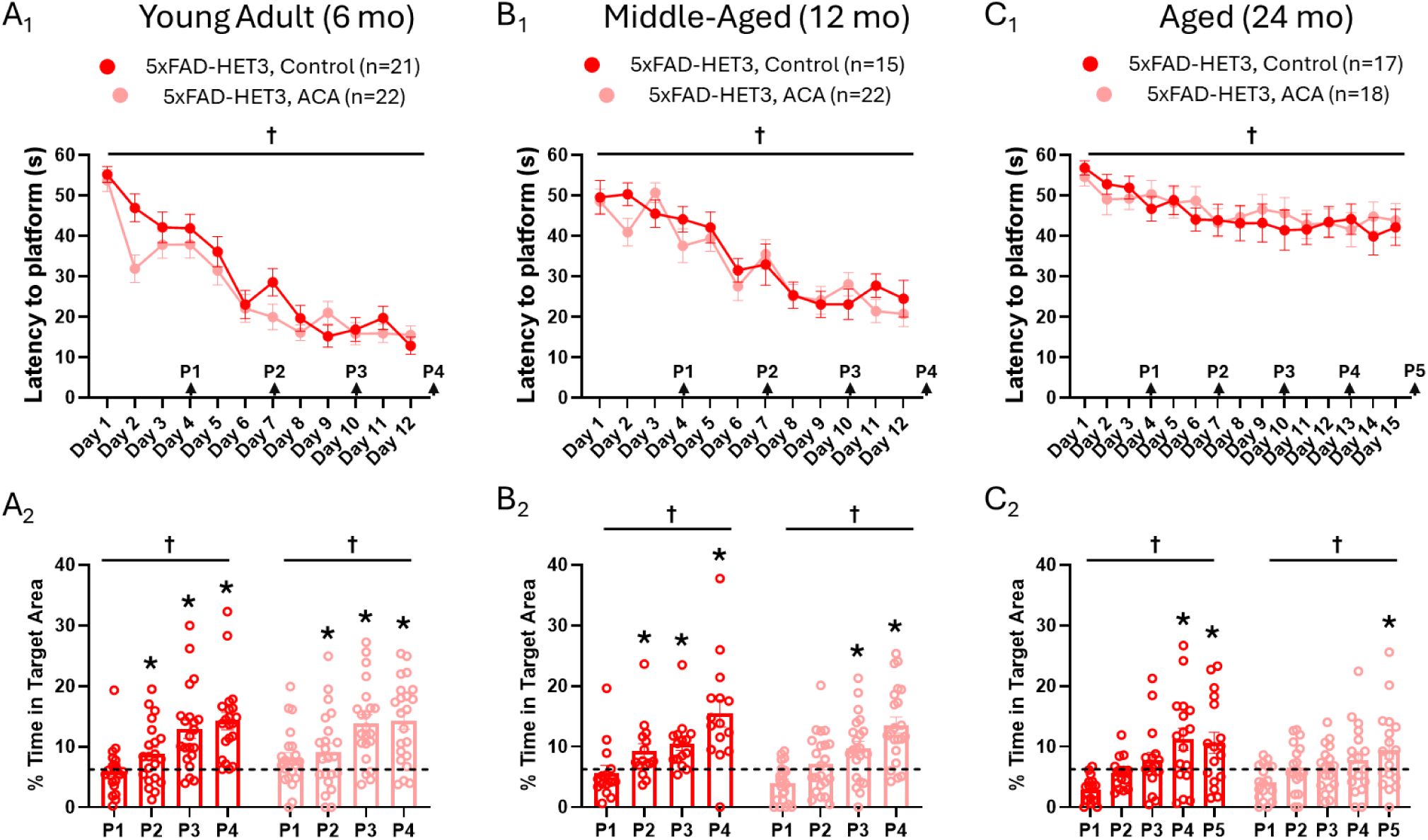
ACA does not ameliorate age-related deficits in spatial learning and memory in 5xFAD-HET3 mice. Separate groups of 5xFAD-HET3 mice were treated with ACA chow (1000 ppm, starting at 4 mo of age; shown in pink) or remained on control diet (“control”, shown in red) and assessed for hippocampus-dependent learning and memory using the MWM at young adult (6 mo), middle-aged (12 mo), or aged (24 mo) timepoints. **(*Top panels, A_1_-C_1_)*** Mice were given 2 trials/day over 12 days of training; the aged group received 3 additional days of training (Days 13 – 15). Both control- and ACA-treated mice in all age groups exhibited a significant reduction in latency to reach the platform over days of training († denotes effect of time, p<0.0001), with no difference in performance between groups at any age (effect of treatment in *A_1_*: young adult, p=0.12; *B_1_*: middle-aged, p=0.47; *C*_1_: aged, p=0.77). See Table 2 for full statistical details. **(*Bottom panels, A_2_-C_2_)*** Four probe trials were interspersed throughout training at the times indicated by the arrows in A_1_-C_1_ (P1 = Probe 1, etc); the aged group received one additional probe trial (P5 = Probe 5). *(A_2_-C_2_)* Both control- and ACA-treated mice in all age groups demonstrated a spatially selective search strategy, as evidenced by a significant increase in the percent time spent searching the target area (circle with 15 cm radius, centered on the trained platform’s location) over successive probe trials († denotes effect of time, p<0.0001), and this performance was not different between treatment groups at any age (effect of treatment in *A_2_*: young adult, p=0.51; *B_2_*: middle-aged, p=0.15; *C_2_*: aged, p=0.47). Further, performance on each probe trial was compared to “chance” (indicated by the dashed line), which is the percent time an animal would be expected to be in the target area “by chance” if they exhibited a random or non-spatially selective search strategy; performance significantly above chance reflects the successful formation of a selective search strategy (* denotes p<0.05). (*A_2_*) In the young adult group, both control- and ACA-treated mice exhibited performance significantly above chance by Probe 2. (*B_2_*) However, in the middle-aged group, while control-treated mice still showed a spatially selective search strategy by Probe 2, this was not demonstrated in ACA-treated mice until Probe 3. (*C_2_*) In the aged group, with formation of a spatially selective search strategy was further delayed until Probe 4 in control-treated mice and Probe 5 in ACA-treated mice. See Tables 3 and 4 for full statistical details.

### 3.6 ACA does not ameliorate age-related deficits in spatial memory in 5xFAD-HET3 mice

Similarly, in probe trials, both control- and ACA-treated mice within each cohort exhibited significant increases in the percent time spent in the target area over the course of the experiment (effect of training in each cohort, p<0.0001) and showed no differences between treatment condition in any age group (effect of treatment in: young adult, p=0.51 ; middle-aged, p=0.15; aged, p=0.47), indicating mice at all ages were able to form a spatial memory for the platform location and that this was not affected by ACA treatment (**Figure 3A_2_-C_2_**; **Table 3A**). Notably, however, when performance was compared between cohorts, both control- and ACA-treated 5xFAD-HET3 mice in the aged group spent significantly less time in the target area than did young adult mice across all probe trials (effect of age, p<0.0001; **Table 3B**), suggesting that there was a significant age-related deficit in memory performance. Consistent with the lack of an effect of ACA within the aged cohort, there was no effect of treatment or interactions of treatment with age between cohorts (effect of: treatment, p=0.89; treatment x age, p=0.37; treatment x training x age, p=0.55; **Table 3B**), indicating that ACA does not improve spatial memory at any age or ameliorate age-related deficits in spatial memory in 5xFAD-HET3 mice.

Age-related deficits in spatial memory were also evidenced by examining individual probe trials to determine when a selective search strategy was exhibited. In the young adult group, both control- and ACA-treated 5xFAD-HET3 mice showed a selective search strategy by Probe 2 (control-treated, p=0.03 and ACA-treated, p=0.04; **Figure 3A_2_**; **Table 4**), but in middle-aged mice, ACA-treated mice did not show a selective search strategy until Probe 3 (p<0.01, while control-treated mice still show performance above chance by Probe 2 (p=0.03; **Figure 3B_2_**; **Table 4**). In the aged group, control-treated mice did not exhibit a selective search strategy until Probe 4 (p=0.02), and aged ACA-treated mice were even further delayed, requiring three additional days of training (Days 13 – 15) to develop a selective search strategy, evident during Probe 5 (p=0.05; **Figure 3C_2_**; **Table 4**). Taken together, these data show that mice in the aged group (regardless of treatment) took the longest to exhibit performance that was statistically above “chance”, reflecting an age-related deficit in spatial memory, and that ACA is not sufficient to ameliorate age-related cognitive decline in an AD-relevant mouse model.

## 4. Discussion

The experiments described here are among the first to evaluate the effect of an anti-aging intervention, which has been shown to extend lifespan, on cognitive function. One previous study examined the effect of another successful lifespan-extending intervention, canagliflozin, on cognitive performance in middle-aged Ntg-HET3 mice and showed that male mice exhibited improved spatial memory (using the Barnes maze) relative to their control-treated counterparts [12]. However, because only one age point was tested, whether canagliflozin was functioning to improve baseline cognitive function (“cognitive enhancer”) or was effectively ameliorating age-related cognitive decline could not be interpreted. Thus, we significantly expand upon this initial study by characterizing cognitive function across the lifespan, including young adult, mid-aged, and aged mice. Our work demonstrates that ACA did not function as cognitive enhancer, as there was no improvement in learning and memory performance in ACA-treated Ntg-HET3 mice in the young adult cohort (at 6 mo of age). Further, our studies show that control-treated Ntg-HET3 mice exhibited age-related cognitive deficits, which were most pronounced in the oldest cohort (at 24 mo), but that both middle-aged (12 mo) and aged (24 mo) ACA-treated Ntg-HET3 mice maintained spatial memory performance equal to that exhibited in young adults. Together, these results demonstrate an ameliorative effect of ACA treatment on “normal” (non-disease related) age-related cognitive decline.

Because aging is the greatest risk factor for the development of AD, we therefore hypothesized that ACA would also be beneficial in reducing cognitive deficits in an AD-relevant mouse model. However, although 5xFAD-HET3 mice also exhibited an age-related progressive worsening of cognitive function, ACA did not improve spatial learning or memory in 5xFAD-HET3 mice at any age, and potentially exacerbated cognitive deficits in the most aged cohort (24 mo). These results underscore the mechanistic divergence between processes underlying age-related cognitive decline in “normal” aging (in the absence of disease) and those that contribute to learning and memory impairments in pathological conditions, like AD. Our findings add to the growing consensus that AD is not a simple extension of normal aging but rather is characterized by distinct pathophysiological mechanisms and differential vulnerability of particular neurological processes. For example, age-related cognitive decline is associated with changes in neuronal function, such as weakening or loss of individual synapses, rather than overt cell death which is characteristic of neurodegenerative disorders like AD [27–29]. Further, the generation, accumulation, and deposition of pathological protein aggregates associated with AD like beta amyloid and hyperphosphorylated tau are generally absent in “normal” aging and likely initiate distinct mechanistic cascades, including a heightened neuroinflammatory response mediated by selective activation of disease-associated microglia, that contribute to the onset and progression of cognitive decline in AD. These differences provide a biological rationale for the selective benefit of acarbose in normal, healthy aging and suggest that interventions focused solely on regulation of peripheral glucose metabolism are unlikely to adequately address mechanistic drivers of AD [27]. Together, these insights highlight not only the limitations of broad-spectrum anti-aging interventions, but also the critical need to develop and refine targeted therapies that specifically address the unique molecular and cellular drivers underlying AD.

Interestingly, while our results suggest that acarbose is not effective in mediating disease-intrinsic deficits in AD, a recent study showed that it may be an effective intervention to address metabolic dysfunction that is exacerbated by AD. When fed a high fat, high sugar (“Western”) diet, 3xTg mice (another AD-relevant mouse model that develops both amyloid and tau pathology [30]) displayed spatial learning and memory deficits (using the Barnes maze) compared to those on a control diet, which were reversed by acarbose [31]. Consistent with our results, acarbose did not significantly affect cognitive function in control diet-fed 3xTg mice, suggesting that acarbose does not directly ameliorate AD-related pathology or cognitive deficits but may instead mitigate metabolic dysfunction that exacerbates AD-related decline, particularly in the context of a Western diet.

An important caveat of our current study is that although we originally balanced our groups with respect to sex, due to unexpectedly high inter-male aggression and attrition, females were overrepresented in our experimental cohorts and limited our statistical power to explicitly evaluate sex-specific effects of ACA on cognitive function.

Interestingly, while ACA treatment (started in young adult mice) significantly extends lifespan in both sexes, the magnitude of the effect is much greater in male HET3 mice compared to females [8, 22]. If the effect of ACA on cognitive function similarly mirrors the observed sex-specific differences in lifespan, our female-biased results may underestimate of the overall efficacy of ACA in promoting cognitive function. Thus, it will be crucial for future studies to rigorously examine the efficacy of ACA in promoting cognitive longevity within well-powered male and female cohorts, enabling robust assessment of both the occurrence and magnitude of sex-specific differences.

While our approach of prioritizing interventions that have demonstrated efficacy in extending lifespan is a logical starting point to identify therapeutics that extend cognitive longevity, it is important to recognize that lifespan and cognition may be mediated, at least in part, by distinct biological pathways and mechanisms. This underscores the importance of testing interventions for effects on cognitive outcomes, even in cases where they have failed to extend lifespan. Interventions that confer meaningful improvements in cognitive longevity, regardless of their lifespan effects, are of great utility and interest in both research settings and clinical applications. Leveraging therapeutic profiles and outcomes associated with distinct interventions provides valuable insights to elucidate the common and distinct mechanistic pathways that govern lifespan and cognitive function. Further, increasing the number of years lived while cognitively intact would not only enhance quality of life and individual autonomy in the aging population, but would also reduce caregiver burden, healthcare cost, and overall impact on society.

Similarly, the geoscience hypothesis suggests that drugs that are effective in slowing aging - by extending lifespan, improving physiological function, or enhancing cognitive longevity - have the potential to ameliorate cognitive deficits specifically associated with age-related diseases, like those associated with AD. However, it is important to continue to assess the efficacy of candidate interventions in the context of AD, and other related dementias, even if they do not demonstrate benefits in “normal aging” populations. While there is likely some mechanistic overlap between normal aging and AD, the pathological impact is also likely to engage distinct disease-related pathways. Therefore, comprehensive testing of these agents in AD models remains warranted, as it may uncover therapeutic benefits that are not evident in healthy aging contexts and help clarify the mechanistic relationships between aging and neurodegenerative diseases.

Finally, in evaluating potential therapeutics, it is essential to assess their effects across a broad spectrum of age- and disease-related paradigms. Not only does this include integrating measures of physical health and lifespan effects with assessments of cognitive longevity, but it should also include evaluations of other physiological domains, such as executive function, attention, working memory, and psychological well-being.

Focusing solely on lifespan extension without considering cognitive and psychological outcomes risks overlooking interventions that may inadvertently exacerbate cognitive decline, diminish well-being, or negatively impact overall quality of life. Therefore, an integrative approach - encompassing not only survival and physical health but also cognitive, emotional, and behavioral function - is necessary to identify treatments that deliver the most impactful improvements in meaningful longevity.

## Acknowledgements

The authors thank members of the Murphy lab and ULAM staff for their assistance with breeding mice, feeding specialized diets, and animal husbandry. We are also grateful to Drs. Stephanie Boas and Catherine Kaczorowski for critical reading and feedback on this manuscript.

